# LC3B conjugation machinery promotes autophagy-independent HIV-1 entry in CD4+ T lymphocytes

**DOI:** 10.1101/2023.07.11.548555

**Authors:** Baptiste Pradel, Maïka S. Deffieu, Véronique Robert-Hebmann, Guilhem Cantaloube, Mathias Faure, Nathalie Chazal, Raphaël Gaudin, Lucile Espert

**Affiliations:** University of Montpellier, 34090 Montpellier, France. CNRS, Institut de Recherche en Infectiologie de Montpellier (IRIM), 34293 Montpellier, France; CIRI, Centre International de Recherche en Infectiologie, Université de Lyon, Inserm U1111, Université Claude Bernard Lyon 1, CNRS, UMR5308, ENS de Lyon, Lyon, France

## Abstract

HIV-1 entry into CD4+ T lymphocytes relies on the viral and cellular membranes’ fusion, leading to viral capsid delivery in the cytoplasm of target cells. The conjugation of ATG8/LC3B protein, process referred to as ATG8ylation and mainly studied in the context of autophagy, occurs transiently in the early stages of the HIV-1 replication cycle in CD4+ T lymphocytes. Despite numerous studies investigating the interplays of HIV-1 with autophagy machinery, the impact of ATG8ylation in the early stages of HIV-1 infection remains unknown. Here we found that HIV-1 exposure leads to the rapid enrichment of LC3B towards the target cell plasma membrane, in close proximity with the incoming viral particles. Furthermore, we demonstrated that ATG8ylation is a key event that facilitates HIV-1 fusion with target CD4+ T cells. Interestingly, this effect is independent of the canonical autophagy pathway as ATG13 silencing does not prevent HIV-1 entry. Together, our results provide an unconventional role of LC3B conjugation subverted by HIV-1 to achieve a critical early step of its replication cycle.

**Teaser:** HIV-1 induces LC3B enrichment towards its target cell entry site and uses the conjugation of this protein to favor its entry step.

## Introduction

Human Immunodeficiency Virus type 1 (HIV-1) entry in CD4+ T Lymphocytes (CD4+ TL) relies on different coordinated steps. Briefly, after its attachment at the surface of target cells, the viral envelope glycoprotein (Env), composed of gp120 and gp41, assembles in trimers at the surface of the viral particle, interacting with their specific receptors CD4 and a co-receptor, CXCR4 or CCR5, depending on the viral tropism. These events lead to the fusion between the viral and cellular membranes allowing the viral capsid delivery into the cytoplasm of the target cell (*1*, *2*). Along its journey towards the nucleus, the viral genome, which is protected by the viral capsid, is retro-transcribed into DNA by the viral Reverse Transcriptase. Once in the nucleus, the viral capsid is uncoated and the retro-transcribed viral genome is inserted in the target cell genome (*3–5*). After this integration step, the viral genome expression produces all the viral components necessary for the formation of new infectious viral particles that will mature after budding from the infected cells (*6*).

In a previous study, we have demonstrated that macroautophagy, evidenced by the ATG8/LC3B conjugation to lipids, is rapidly and transiently induced upon HIV-1 infection of CD4+ TL (*7–9*). Macroautophagy, hereafter referred simply to as autophagy, is a major lysosomal degradation and recycling pathway involved in many cellular processes (*10–12*). Notably, it is a protective mechanism against infections able to directly degrade pathogenic components or by regulating innate and adaptative immunities (*13*, *14*). This process relies on the formation of double membrane vesicles, called the autophagosomes, which sequester cytoplasmic components that are degraded after their fusion with lysosomes. The autophagy initiation is tightly regulated by several signaling complexes such as the ULK1/2-FIP200- ATG13 complex and the class III phosphatidylinositol 3-kinase/BECLIN 1 (PI3KC3/BECN1) complex (*13*). Autophagy is commonly monitored by following the conjugation of the ATG8/LC3B family members to a lipid, the phosphatidylethanolamine (PE) because the ATG8/LC3B-PE conjugate is anchored in the autophagosomal membranes and is degraded together with the substrates in the lysosome, making it a good marker of the autophagic flux (*14*). Even if ATG (AuTophaGy-related) proteins have been identified for their involvement in autophagy, it is now clear that they also exert independent functions (*15*, *16*). In particular, the conjugation of the ATG8/LC3B family members to lipids, recently named “ATG8ylation” by analogy with ubiquitination, has been proposed to be a general mechanism involved in the maintenance of membrane homeostasis during remodeling (*17*, *18*). Interestingly, it has been shown that the conjugation of ATG8 proteins is not restricted to PE and that this protein can also be conjugated to phosphatidylserines (PS) (*19*, *20*). ATG8ylation involves an E1-like ATG7 protein, two E2-like ATG3 and ATG10 proteins and finally the E3-like complex ATG5−ATG12-ATG16L1. The recruitment of these ubiquitin-like conjugation systems to membranes requires the upstream production of PI3P (Phosphatidylinositol-3-phosphate) by the PI3KC3/BECN1 complex (*21*). It is worth noting that, so far, the ULK1/2-FIP200-ATG13 complex is only required for canonical autophagy, allowing to make a clear distinction with others ATG8ylation-mediated processes (*22*).

During their replication, viruses have to face different cellular and immunological barriers in order to achieve a successful replication cycle. As a consequence, along evolution, viruses have adapted to either block or hijack cellular processes, in particular the autophagic machinery, to their own benefit (*23*). Indeed, autophagy is rapidly activated upon viral infections and contribute to viral sensing allowing a tight control of the infection. For example, the engagement of the Measle virus (Edmonston strain) cell surface receptor CD46 induces autophagy in the early step of infection *via* an indirect association with BECN1 and Herpesviruses are able to stimulate autophagy in the early steps of infection before inhibiting the autophagic maturation step to avoid their degradation (*24–26*). Nothing is currently known about the interplay between the autophagic machinery and HIV-1 during the early steps of CD4 TL infection but it has been demonstrated that the downregulation of several ATGs favors HIV-1 infection in these cells (*27*). This effect has also been observed through a functional genomic screen showing the pro-HIV-1 role of some ATG proteins, in particular ATG7 and ATG12, which are known to be involved in the conjugation of ATG8 proteins (*28*). Consequently, the aim of our study was to determine the consequence of the early induction of ATG8/LC3B (hereafter only referred as LC3B) conjugation, observed during HIV-1 CD4+ TL, on viral replication. The results presented here show that LC3B is enriched at the plasma membrane (PM) of the target cells, in close proximity of the HIV-1 entry sites. Moreover, we show that the conjugation machinery of LC3B favors the HIV-1 entry step in CD4+ TL independently of canonical autophagy. Altogether, our results bring unprecedent information for the understanding of the cellular mechanism used by HIV-1 to promote the first step of its replication cycle.

## Results

### PI3P production favors HIV-1 replication in CD4+ T lymphocytes

The ATG8 protein LC3B is conjugated to lipids during the early steps of HIV-1 replication in CD4+ TL suggesting that HIV-1-induced autophagy could influence the viral replication (*7*). Moreover, several ATG proteins were shown to favor HIV-1 infection (*27*, *28*). To evaluate the role of the autophagy machinery during HIV-1 replication, we inhibited the production of PI3P, a phospholipid required for the recruitment of the ATG8ylation machinery, by using a specific inhibitor of the PI3KC3/BECN1 complex (PIK-III (*29*)). We pre-treated CD4+ T cells, either a T cell line (HuT 78) or primary CD4+ TL purified from the blood of healthy donors, with PIK-III during 3 h before infecting them with purified HIV-1 particles. After 1 h of contact with the viruses, target cells were extensively washed and HIV-1 replication was monitored 16 h later by quantifying the intracellular (cell lysates) and extracellular (culture supernatants) HIV-1 capsid protein p24 by ELISA. The results showed that the inhibition of PI3P production leads to a significant decrease in both the intracellular and the extracellular p24 levels (Fig. 1A).

**Figure 1.**
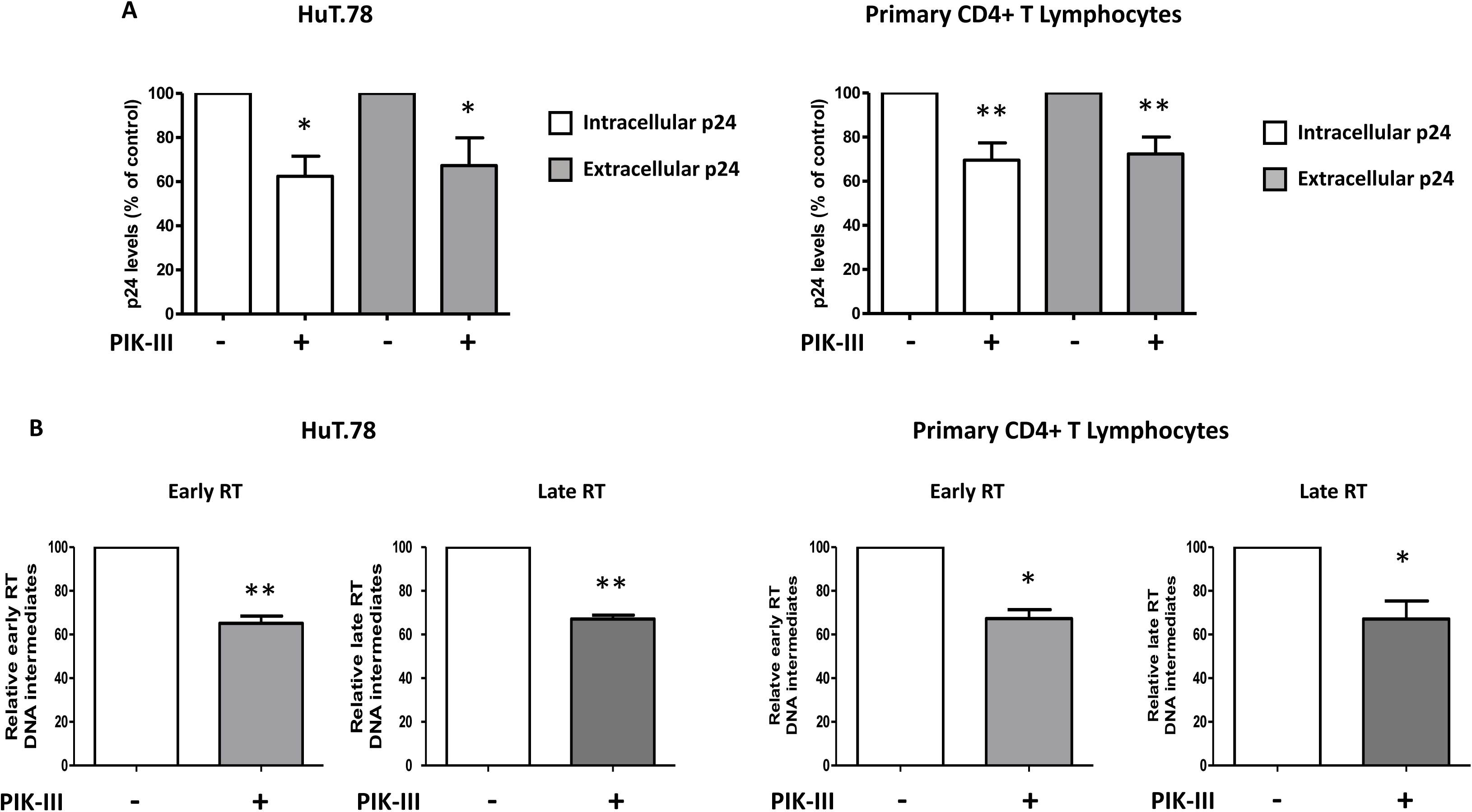

After viral capsid delivery into target cells, the HIV-1 RNA genome is retro-transcribed into DNA before being integrated in the cellular genome. To analyze the impact of PI3P production on this step of the viral replication cycle, we quantified both the early and late reverse transcribed viral products in untreated- and PIKIII-pretreated HIV-1-infected cells. As presented in the Fig. 1B, we observed that addition of PIK-III decreased both viral reverse transcribed DNA products.

Altogether, by using two complementary approaches, we showed that PI3KC3/BECN1 complex is required for HIV-1 replication in CD4+ TL, indicating that the autophagy machinery may play an important role in the early stages of the HIV-1 life cycle.

### LC3B puncta are enriched toward the target cell PM upon HIV-1-like particles exposure

As we showed above, both p24 and viral reverse transcribed DNA products level are affected by the PIK-III pre-treatment suggesting that the autophagy machinery plays an important role in the very first steps of HIV-1 replication. In order to monitor the early viro-induced autophagic events at the single cell level, we performed live-cell imaging to analyze the dynamics of LC3B puncta formation upon virus exposure, a reliable readout to monitor the activation of the autophagy machinery. To avoid overexpression artifacts, we used a CRISPR/Cas9-based approach to generate HEK 293T cells expressing endogenous levels of a fluorescently tagged LC3B (GFP-LC3B^edit^). To this aim, a sequence coding for the GFP protein was inserted upstream of the gene coding for LC3B (Fig. 2A). After two rounds of cell sorting, we obtained a partially edited HEK GFP-LC3B^edit^ cell line expressing endogenous levels of GFP-tagged LC3B (Fig. S1). To characterize the functionality of our model, we starved HEK GFP-LC3B^edit^ cells using nutrient-free medium (EBSS), a well-known inducer of autophagy, during 4 h in the presence of Bafilomycin A1 (BafA1), which is known to impair the autophagy flux. Microscopic analyses of LC3B puncta showed that EBSS+BafA1 co-treatment expectedly resulted in an accumulation of autophagosomes. Moreover, all the LC3B puncta, revealed by the anti-LC3B antibody, colocalized with the GFP-LC3B^edit^ puncta, further validating our model for subsequent analyses (Fig. 2B). This cell line was then transduced to express the HIV-1 receptors in order to make them permissive to HIV-1 infection. Hence, we generated a novel HEK GFP-LC3B^edit^.CD4.X4.R5 cell line, which expresses CD4, CXCR4 and CCR5. HEK GFP.LC3B^edit^.X4 cells, which express only CXCR4, was also obtained for the need of subsequent experiments (Fig. 2C).

**Figure 2.**
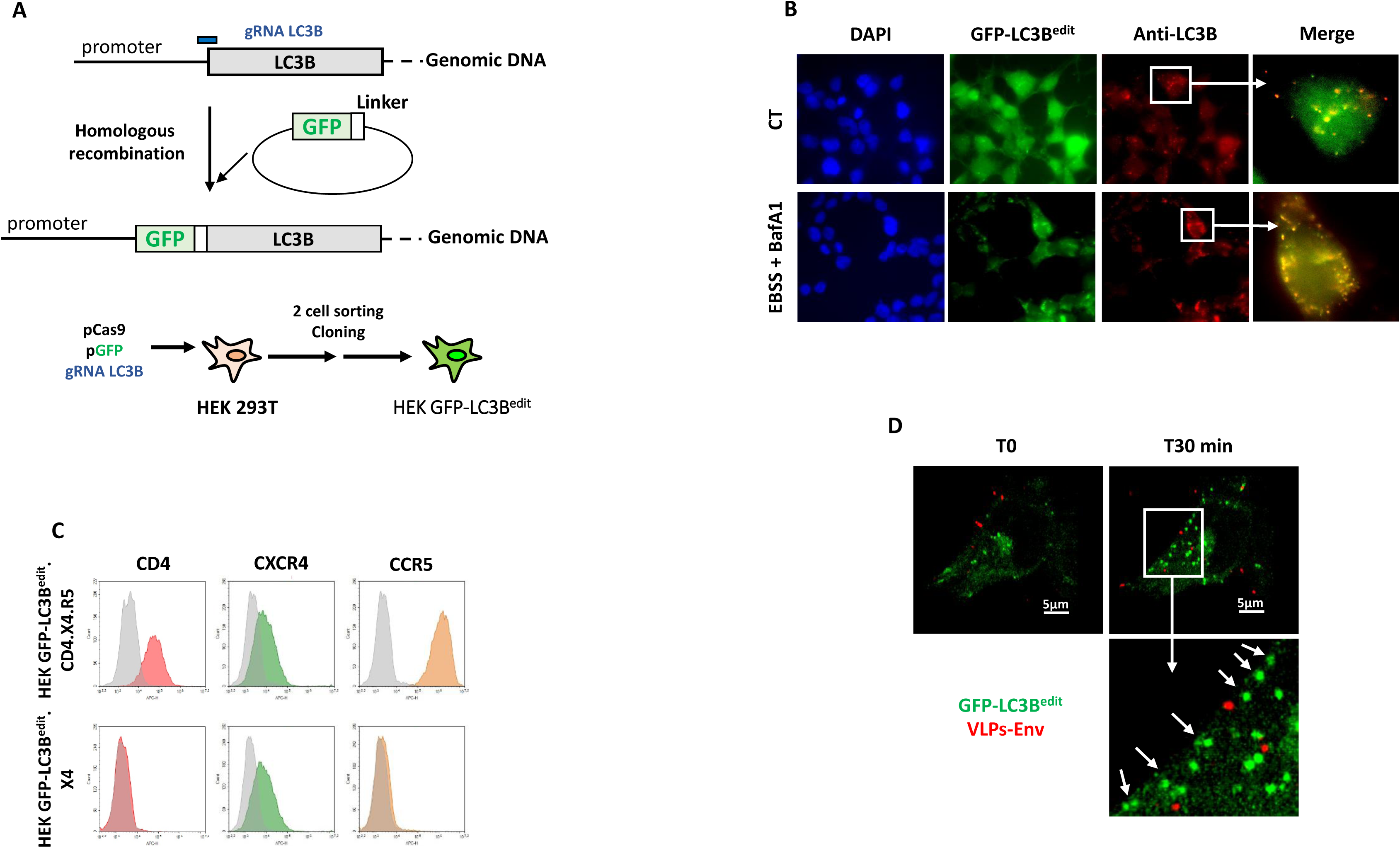

In parallel, to study the trafficking of the virus, we produced fluorescent virus-like particles (VLPs) composed of a fluorescent HIV-1 core (mCherry-tagged Gag protein) surrounded by an envelope expressing the HIV-1 Env at its surface (VLPs-Env, NL4.3 strain). These VLPs- Env were used to mimic an HIV-1 infection of the HEK GFP-LC3B^edit^.CD4.X4.R5 cells and to follow the interplay between viral particles and GFP-LC3^edit^ puncta dynamics by live-cell imaging. As shown in Fig. 2D and in Movie S1, this strategy allowed us to observe an increase in the number of GFP-LC3B^edit^ puncta in VLPs-Env-exposed cells overtime. Strikingly, we noted an enrichment of these GFP-LC3B^edit^ puncta towards the target cell PM, which is clearly observable after 30 min of VLPs-Env exposure.

### The presence of CD4 is required to observe the LC3B enrichment towards the target cell PM upon HIV-1-like particles exposure

To quantify the enrichment of GFP-LC3B^edit^ puncta, at the target cell PM, induced by exposure to VLPs-Env, we stained the target cells with the SirActin dye. This dye, which stains the subcortical polymerized actin, allowed us to visualize the cell border and, then, to measure the distance between each GFP-LC3B^edit^ puncta inside the target cell from the PM. The data presented in the Fig. 3A showed a significant decrease in this distance upon VLPs- Env exposure, confirming the viral-induced enrichment of GFP-LC3B^edit^ puncta at this site of the target cells. Upon interaction with its target cells, the gp120 subunit of Env interacts first with CD4, leading to a conformational change allowing its subsequent interaction with the co-receptor. This last interaction allows the gp41 subunit of Env to insert in the target cell PM to achieve the fusion between the viral and the target cell membranes, ultimately leading to the viral capsid entry (*2*, *30*). In order to determine which of these events is involved in the viro-induced enrichment of LC3B puncta towards the target cell PM, we first used AMD3100, an antagonist of CXCR4 that blocks its interaction with gp120. Analyses of the distance between GFP-LC3B^edit^ puncta and the Sir-Actin staining revealed that the addition of AMD3100 did not prevent the viro-induced enrichment of GFP-LC3B^edit^ puncta towards the target cell PM (Fig. 3B). To test the involvement of CD4, the same experiment was done in edited target cells that do not express this receptor (HEK GFP-LC3B^edit^.X4, Fig. 2C). In these conditions, the viro-induced enrichment of LC3B puncta towards the PM was abolished (Fig. 3C). This result is likely due to the lack of CD4 rather than CCR5 since the VLPs-Env do not use CCR5 for entry. Consistently, the same result was obtained in target cells that do not express CD4 and are treated with AMD3100 (Fig. S2). Altogether, these results strongly suggest that the gp120-CD4 interaction is involved in the enrichment of LC3B puncta towards the target cell PM.

**Figure 3.**
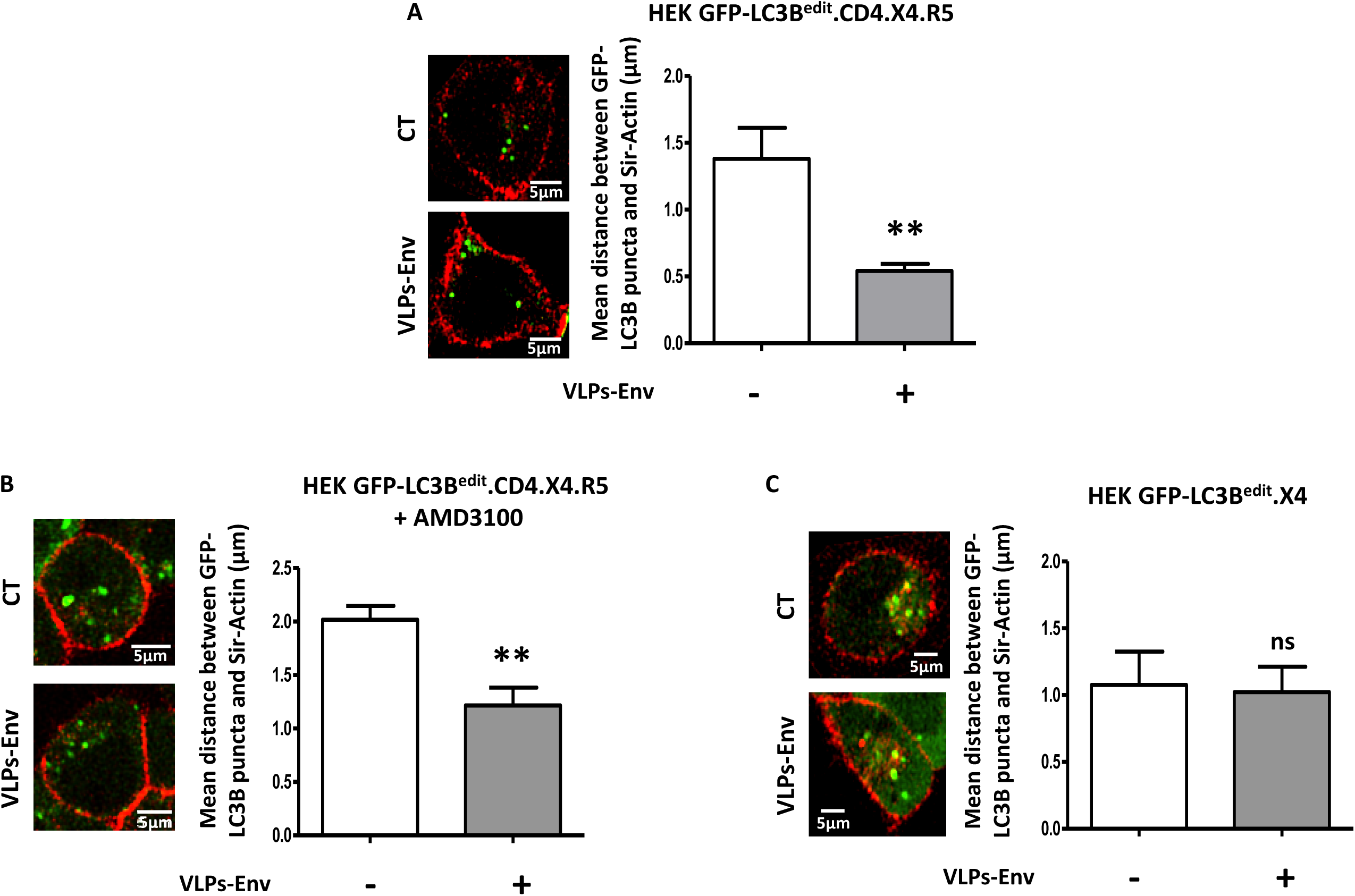

### HIV-1 exposure leads to an enrichment of LC3B puncta towards the virus entry sites

To go deeper in the characterization of the VLPs-Env-induced enrichment of LC3B puncta towards the target cell PM and to determine if this enrichment specifically occurs near the viral particle entry sites, we tracked the fluorescent-tagged VLPs-Env together with GFP- LC3B^edit^ puncta by live-cell imaging. To achieve this goal, we delimited a zone, by drawing a sphere of 1.5 µm^3^ at the surface of the target cell before the arrival of the VLPs-Env. We analyzed the mean number of GFP-LC3B^edit^ puncta in this zone, before, during and after the presence of the VLPs-Env at the cell surface, by tracking VLPs-Env particles over time. As shown in Fig. 4A and in the associated Movie S2, no GFP-LC3B^edit^ puncta were observed at the cell surface in absence of the VLPs-Env in the delimited zone. Then, a concomitance between PM-proximal GFP-LC3B^edit^ puncta appearance and the VLPs-Env fluorescence intensity appeared and persisted all along the presence of the viral particles at the target cell surface. Finally, both the VLPs-Env fluorescence intensity and the GFP-LC3B^edit^ puncta disappeared after the viral particle entry. This result was quantified by tracking several fluorescent viral particles. As the experiments presented in Fig. 4A were done using VLPs- Env, we next analyzed the behavior of GFP-LC3B^edit^ puncta after exposure to fluorescent wild-type viruses. To do that, we infected the edited cells with X4 or R5-tropic fluorescent wild-type HIV-1 and we applied the same protocol as previously. As shown in Fig. 4B and 4C, both X4-tropic HIV-1 and R5-tropic HIV-1 triggered transient PM-proximal GFP-LC3B^edit^ puncta appearance near the viruses (associated Movie S3 and S4, respectively). Altogether, these results indicated that LC3B puncta are preferentially enriched towards the virus entry sites independently of the co-receptor used.

**Figure 4.**
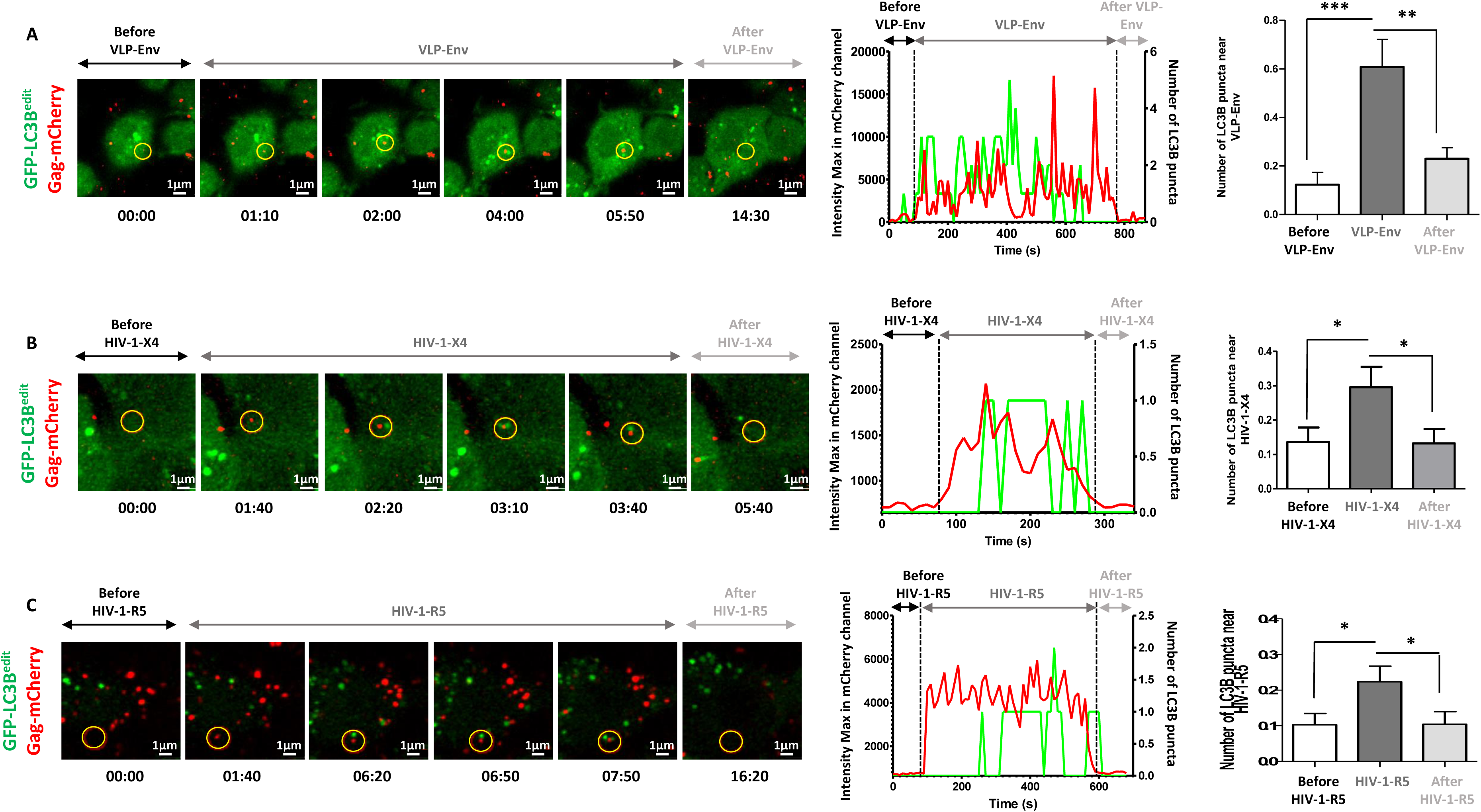

### LC3B conjugation favors HIV-1 entry in target CD4+ T cells

Our data suggest that LC3B could play an important role in the HIV-1 entry step. To test this hypothesis, we first analyzed the impact of PI3P production on this step of the viral replication cycle. To this aim, we used a standard method, which is the BlaM (Beta-Lactamase)-Vpr assay (*31*). It consists in producing virions having incorporated the BlaM fused to the HIV-1 Vpr protein, which is enclosed in the viral particles. These virions were purified and used to infect CD4+ T cells stained with the fluorescent CCF2, which is the substrate of BlaM. Consequently, upon HIV-1 entry, the BlaM-Vpr fusion protein, released in the target cell cytosol, cleaved the CCF2 substrate. As CCF2 emits a different fluorescence upon cleavage, it allows the quantification of HIV-1 entry, occurring after the fusion between viral and cellular membranes, by flow cytometry. As shown in the Fig. 5A, PIK-III pretreatment led to a significant decrease in HIV-1 entry, both in HuT 78 cells and in primary CD4+ TL, demonstrating that PI3P production is involved in HIV-1 entry in target CD4+ T cells. Importantly, we validated that the drug did not affect the expression level of the HIV-1 receptor and co-receptor, CD4 and CXCR4, at the cell surface (Fig.S3). As PI3P is required for the recruitment of the LC3B conjugation machinery and because we previously observed a rapid LC3B lipidation upon HIV-1 infection of CD4+ T cells (*7*), we next investigated the role of this process during the HIV-1 entry step. To achieve this goal, we transduced CD4+ T cells with shRNA allowing the downregulation of the expression of both BECN1, which is involved in PI3P production, and ATG5, which is essential for the conjugation of LC3B to lipids. Then, both shRNAs-transduced cells were infected with BlaM-VprR-containing HIV-1 and the level of viral entry was analyzed by flow cytometry as above. The results obtained showed that both the downregulation of BECN1 and ATG5 expression leads to a decreased HIV-1 entry in target cells (Fig. 5B) arguing for a role of LC3B conjugation in the observed effect. In order to confirm the critical role of LC3B lipidation in HIV-1 entry into target cells, we did the same BlaM-Vpr assays in CD4+ T cells over-expressing LC3B or a LC3B-mutated form unable to be lipidated (LC3B G120A). As shown in Fig. 5C, LC3B over-expression increased HIV-1 entry in target cells whereas the mutated LC3B, incapable of lipidation, had no effect on this first step of the viral life. These data suggest that HIV-1 particle fusion with target cell membranes is dependent on the intracellular lipidation function of LC3B.

**Figure 5.**
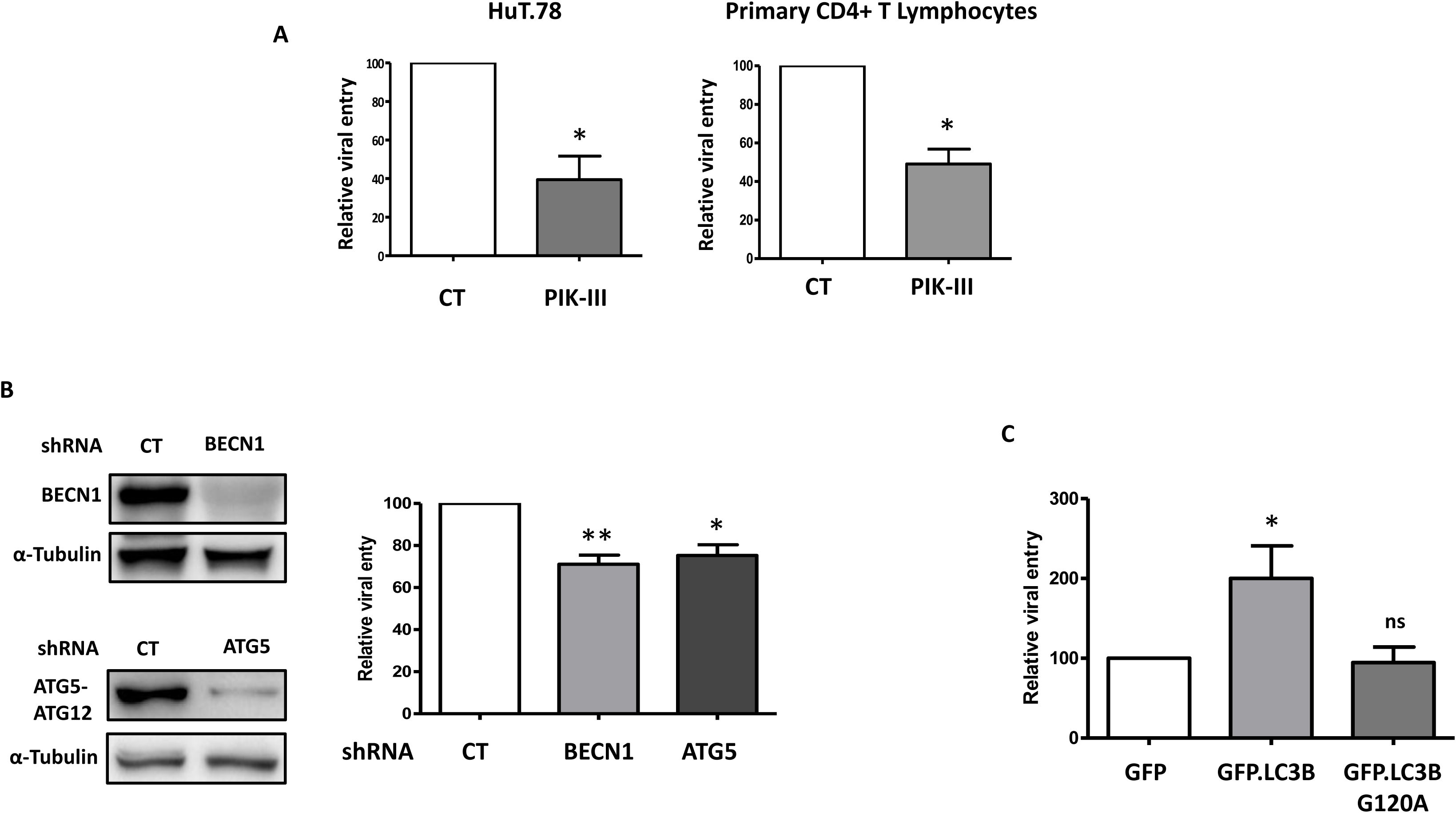

### LC3B conjugation favors HIV-1 entry in CD4+ T cells independently of the canonical autophagy pathway

Nowadays, ATG8ylation is well-known to be involved in several processes besides the canonical autophagy pathway and it is proposed to be a general membrane stress and remodeling response (*32*). In order to determine if the effect of ATG8ylation on HIV-1 entry is due to canonical autophagy or an independent process, we analyzed this step using a cocktail of lysosomal inhibitors targeting cathepsins B and D (E64d + Pepstatin A). As shown in Fig. 6A, inhibition of lysosomal degradation led to an increase in HIV-1 entry in CD4+ T cells. These data are in accordance with previous reports showing that inhibition of endosome acidification with BafA1, a drug also known to inhibit autophagic degradation, enhances entry by fusion at the PM (*33*). To specifically address the role of canonical autophagy during HIV- 1 entry, we transduced shRNAs to downregulate ATG13 expression and we tested the effect of this downregulation on HIV-1 entry using the BlaM-Vpr assay. Indeed, ATG13, which is part of the ULK1/FIP200/ATG13 signaling complex, is known to be only involved in the initiation of canonical autophagy allowing to discriminate between the role of LC3B conjugation in canonical autophagy or in an independent process. Interestingly, we observed that the downregulation of ATG13 led to an increase of HIV-1 fusion to target cells (Fig. 6B), an opposite effect than the one observed with shRNA targeting BECN1 or ATG5. One hypothesis that could explain this phenotype is that releasing the autophagic machinery components from their role in canonical autophagy would allow the ATG8ylation actors to be more available to act on viral entry. Taken together, these results show that the conjugation of LC3B to lipids participates in HIV-1 entry in target cells independently of an autophagy-mediated degradation.

**Figure 6.**
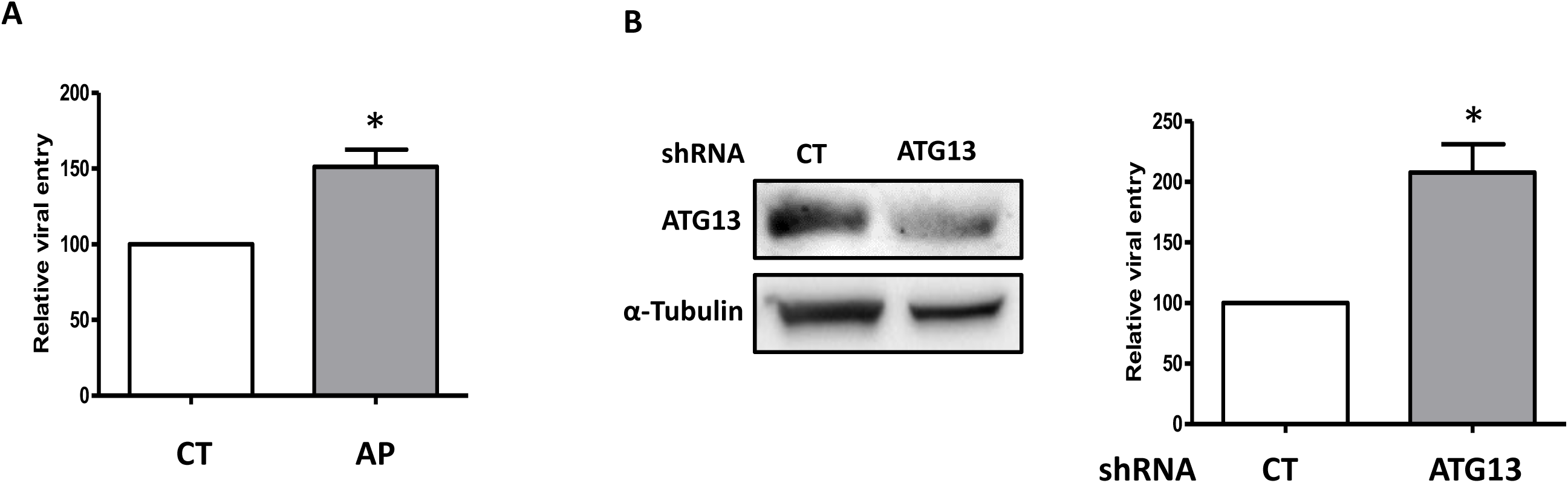

## Discussion

HIV-1 entry in its target cells is the first step of a successful replication cycle. Our results show that, even if the mechanisms driving HIV-1 entry has been extensively studied since the discovery of this virus forty years ago, new technical approaches can bring additional insights in this highly dynamic process. Indeed, we demonstrate, for the first time, that ATG8ylation can influence the entry step of a virus in its target cells. By using live-cell imaging, we showed that the LC3B protein is enriched towards the target cell PM upon HIV-1 exposure, particularly near the virus entry sites. In particular, the conjugation of this protein to lipids, process recently named ATG8ylation, favors HIV-1 entry independently of the canonical autophagy process. Our results are in accordance with a previous study showing that peptides derived from the ATG protein BECN1 and BECN2 (Tat-BECN1 and Tat-BECN2, respectively) favor lentivirus entry into target cells. These peptides, which have been first developed as canonical autophagy inducers (*34*), exert their pro-entry effect at doses that do not induce canonical autophagy but leads to LC3B conjugation (*35*).

Interestingly, our results showed an increase in HIV-1 entry after the addition of lysosomal cathepsins inhibitors (Fig. 6A), indicating that the lysosomal degradation could restrict HIV-1 entry. However, it is well known that using these inhibitors leads to LC3B-positive membranes accumulation (*14*). Thus, lysosomal inhibitors treatment could increase the quantity of LC3B-conjugated membranes that would be available for HIV-1 fusion process. However, we cannot exclude that several lipidation sites could be involved. Indeed, our data do not allow to determine if the LC3B lipidation occurs locally, directly on lipids in the vinicity of the PM, or on lipids from intracellular membranes that would subsequently be redirected towards viral entry sites. In addition, it would be interesting to determine if some proteins from the endoplasmic reticulum (ER) are localized at the PM during HIV-1 entry since it has been shown that LC3B lipidation could be driven by a local synthesis of PI3P at the level of ER-PM contact sites (*36*, *37*). Concerning the PM lipid composition needed for HIV-1 entry, despite numerous discrepancies, several studies pointed out the roles of cholesterol and ordered lipid domains in HIV-1 cellular entry and indicate that viral fusion occurs preferentially through cholesterol-rich heterogeneous membranes and not through uniformly fluid membranes (*38*, *39*). Afterwards, it has been shown that ordered and disordered lipid domains coexistence is required for a successful fusion step (*40*). Thus, it would be interesting to analyze if LC3B lipidation could influence locally the lipids composition and organization at the PM and modify its fluidity and stiffness.

At the molecular level, the interaction of HIV-1 Env with CD4 appears to be required for the LC3B enrichment at the target cell PM suggesting a role of signaling events downstream of this receptor. In this context, it has been reported that the Rac1 signaling complex, involved in actin cytoskeleton remodeling and known to be activated by HIV-1 Env, promotes HIV-1 entry into target cells by acting on membrane fusion (*41*). In parallel, another study showed that infection by *Campylobacter jejuni* leads to a Rac1-dependent recruitment of LC3B at its host cell entry site, and that the conjugation of this protein favors bacterial invasion (*42*). These results suggest that the LC3B enrichment towards the PM could be a more general pathogen-induced phenomenon, independent of a specific cellular receptor. Consequently, other cellular partners shall be sought for. In particular, it is well known that adhesion molecules expressed at the cell surface, such as integrins, are the first cellular components that interact with incoming pathogens. The role of these proteins has been largely studied in the context of HIV-1 infection and are known to facilitate the recognition between HIV-1 Env and its specific receptors (*43–46*). It is worth noting that a recent study has shown that integrins harbor a LIR motif (LC3-Interacting Region) in their cytoplasmic part (*47*). Thus, one could hypothesize that integrins would be involved in the recruitment of LC3B towards HIV-1 entry sites. In this context, as the Env/integrin interaction is transitory, the subsequent Env/CD4 interaction would be required to maintain LC3B at the PM enough time in order to favor viral and cellular membranes fusion and could explain our results showing that Env/CD4 interaction is required to observe the LC3B enrichment towards the PM. In addition, integrin signaling is known to trigger a calcium response needed for cellular activation (*48*, *49*). Moreover, a Ca^2+^-dependent signaling pathway is triggered after interaction between Env with its receptors (*41*, *44*, *50–53*) and promote the Env-mediated membrane fusion by inducing a transient redistribution of PS from the inner leaflet to the outer leaflet of the PM (*54*). As Ca^2+^ signaling and autophagic membranes synthesis has been intimately connected in several studies (*55*, *56*), it could be interesting to determine if the LC3B conjugation, to PE or PS, could be dependent on Ca^2+^ upon HIV-1 exposure and, by this way, could regulate the virus-host cell fusion events.

ATG8 is unique in yeast but seven homologs have been identified in mammals. These proteins are classically divided in two families, which are the LC3 family, composed of LC3A, LC3B, LC3B2 and LC3C, and the GABARAP (γ-aminobutyric acid receptor-associated protein) family composed of GABARAP, GABARAPL2 and GABARAP L2. Several reports demonstrated a specific role of ATG8 proteins in mediating membrane fusion. Indeed, *in vitro* studies demonstrated that the fusion between two liposomes is favored by the expression of the yeast ATG8 at their surface. The authors also observed an enrichment of these ATG8 at the junction between the liposomes (*57*). Accordingly, another study, using the mammals ATG8s LC3B and GABARAPL2, showed that liposomes-bound ATG8s can induce their fusion (*58*). Recently, it has been demonstrated that both the yeast ATG8 and LC3B can trigger membrane perturbations suggesting a role of these proteins in membrane remodeling (*59*). Considering the amount of data in the literature regarding the contribution of the ATG8 proteins on membrane fusion, it would be important to decipher their specific actions, individually or in cooperation with each other’s, on fusion mechanisms taking place at the PM during HIV-1 entry.

To our knowledge, despite several examples described of ATG8ylation in cellular secretion, such as unconventional secretion or viral release (*60–65*), the present study is the first to identify such a role during internalization of exogenous cargoes. This discovery could have wide repercussions on the characterization of different cell biology processes taking place at the PM, such as, for example, extracellular vesicles entry in cells. In the case of HIV-1 infection, it would be critical to identify the cellular and the viral partners of LC3B in the context of viral entry as the discovery of new actors in this process could lead to the development of new combined therapies in the future.

## Materials and Methods

### Cells

HuT 78 cells were cultured and maintained in RPMI 1640 medium supplemented in 10% FCS, 1% penicillin/streptomycin. HEK 293T cells were cultured and maintained in cultured in Dulbecco’s modified Eagle’s medium (DMEM) supplemented with antibiotics and 10% fetal calf serum (FCS). Primary CD4+ T lymphocytes were isolated from peripheral blood mononuclear cells and purified by negative selection using the CD4+ T cell isolation kit (Miltenyi Biotec, 130-096-533). Cells were phenotypically analyzed and activated using anti- CD3/antiCD28 coated plates (Ref BE0001-2 and Ref BE0248, respectively, BioXCell) followed by IL-2 treatment (100U/mL Miltenyi Biotec 130-097-746). All experiments involving primary cells were performed at least 4 days post-activation.

Obtention of edited cells (HEK GFP.LC3B^edit^.X4 and HEK GFP-LC3B^edit^.CD4.X4.R5): The HEK.GFP.LC3B^edit^.X4 was generated according the method in (*66*). Briefly, we transfected HEK 293T cells with a combination of a donor plasmid (p-GFP: MAP1LC3B-mEGFP gifted from Allen Institute for Cell Science - Addgene Ref 101783), a purified PCR product coding for the gRNA-tracrLC3B and a plasmid coding for Streptococcus pyogenes Cas9 (pCas9) using JetPrime (Polyplus Transfection). 7 days post-transfection, cells were sorted to be enriched using the cell sorter Aria IIU (Becton Dickinson). A second cell sorting was performed to obtain several clones selected according to the GFP expression levels and a single clone (HEK.GFP.LC3B^edit^.X4) was selected. A genomic DNA extraction was performed on this clone using the Quick DNA MicroPrep kit (Zymo Research Ref D3020) using manufacturer’s instructions and a subsequent PCR analysis was done using primers amplifying both the LC3B gene and the GFP-edited copy of this gene using Taq Platinium Polymerase Kit (Invitrogen, Ref 10966034) (primers sequence: 5’- GCGGGCTGAGGAGATACAAG-3’ and 5’-TCCGAGAGGTCCGCAGG-3’). PCR products were run on a 1% agarose gel and imaged with SYBR Safe DNA gel stain (invitrogen Ref S33102) using Axygen Gel Documentation System. The edited cells were then transduced with lentivectors expressing CD4 and CCR5 in order to obtain the HIV-1 permissive cells named HEK GFP-LC3B^edit^.CD4.X4.R5.

### Reagents and antibodies

PIKIII was purchased from Euromedex (Ref SE-S7683); AMD3100 (Ref A5602), Baf A1 (Ref B1793), Pepstatin A (Ref 77170) and E64d (E8640) were purchased from Sigma-Aldrich. Sir-Actin was purchased from Tebu-Bio (Ref 251SC001). The anti-LC3B (Ref L7543) and the anti-ATG5 (Ref A0856) were purchased from Sigma-Aldrich. The anti– BECN1 Ab (Ref 11306-1-AP) was purchased from Proteintech. The anti-ATG13 (Ref 13468) was purchased from Cell Signaling. The anti-α-Tubulin was purchased from Sigma-Aldrich (T5168). The anti-CD4 antibody (clone 13B8.2) was purchased from Beckman, the anti- CXCR4 antibody (clone MAB173) from R&D systems and the anti-CCR5 (2D7) from Becton Dickinson.

### Viruses

*X4-tropic viruses:* HEK 293T cells were transfected with pNL4-3 using PEImax (Clinisciences). *Fluorescent X4-tropic viruses:* HEK 293T cells were co-transfected with pNL4.3 and pGag.imCherry. *Fluorescent R5-tropic viruses:* HEK 293T cells transfected with the pNL4.3.Gag-imCherry molecular clone, which carries an R5-tropic envelope thanks to a mutated gp120 V3-loop (*67*). *VLPs-Env:* HEK 293T cells were co-transfected with pCDNA3.Env (from NL4.3), psPAX2, pGagmCherry and pLenti-GFP. *X4-tropic BlaM-Vpr viruses*: HEK 293T cells were co-transfected with pNL4.3 and pBlaM-Vpr. Viral particles were concentrated by ultracentrifugation on a PBS-Sucrose 25% cushion after the collection of the culture supernatants 48h post-transfection. Quantification of viral particle production was performed with a p24 enzyme-linked immunosorbent assay (ELISA) from Innogenetics (Innotest HIV antigen monoclonal Ab [MAb]) according to the manufacturer’s instructions.

### Lentiviruses production and transduction

HEK 293T cells were co-transfected with pMD2.G (expressing VSV-G envelope protein), pPAX2 and pLKO.1-puro shRNAs expressing plasmids (MISSION, Sigma-Aldrich). Produced lentiviruses were concentrated and quantified as previously. Cells were transduced with shRNAs-containing lentiviral particles for 48h and, then selected using puromycin during 10 days before being used for experiments.

To obtain the GFP constructs-expressing lentiviruses, HEK 293T cells were co-transfected with pMD2.G, pPAX2 and pNaldini-eGFP or pNaldini-eGFP-LC3B. The pNaldini-eGFP- LC3B.G120A was obtained by directed mutagenesis using the Q5 site directed mutagenesis kit (Ref: E0554S, New England Biolabs). Produced lentiviruses were concentrated and quantified as previously. Target CD4+ T cells were transduced with GFP or GFP-LC3B or GFP-LC3B.G120A lentiviral particles (ratio of 4 :1 between GFP and GFP-LC3B or GFP- LC3B.G120A constructs to obtain the same percentage of transduction) during 3 days before experiments.

### Analysis of HIV-1 reverse transcribed products by qPCR

HuT 78 cells or primary CD4+ T cells were infected with DNase I-pretreated wt-X4-tropic viruses for 16h. DNA from infected cells was extracted using the QIAamp DNA-minikit (Qiagen) according to manufacturer’s instructions. Quantitative PCR of early and late retrotranscribed products was done using the TaqMan Universal Master mix II no UNG (Ref 4440040, ThermoFisher) and the quantity normalized to PGBD loading control. Analysis was performed with LightCycler 480 with the method of 2ΔΔCT.

### Western Blot

Cell pellets were lysed in 2x Laemli Buffer and heated 10 minutes at 95°C. Samples were loaded on 12% Prosieve 50 gels (Lonza, LON50618) and transferred to polyvinylidene (PVDF) membranes. After a blocking step in TBS containing 5% casein (Sigma-Aldrich, C7078) and 0.05% Tween 20 (Sigma-Aldrich, P9416) for 1 h at room temperature, blots were incubated overnight at 4°C with the primary Ab in the blocking buffer. After 3 washes with TBS+0.05% Tween 20, the blots were incubated for 1 h at room temperature with peroxidase-coupled antiserum (Sigma-Aldrich, goat anti-rabbit and goat anti-mouse, A0545 and A2304, respectively) diluted in blocking buffer. After further washes, the immune complexes were revealed by ECL (Clarity western ECL Substrate; Bio-Rad, 170–5061) and analyzed with a ChemiDoc camera (Bio-Rad). Quantification of protein expression was performed using the Image lab Software (Bio-Rad). Data were normalized with reference to the densitometry analysis of α-tubulin.

### BlaM-Vpr assay

Target cells were infected with X4-tropic BlaM-Vpr viruses at a MOI of 1 for 3h. After a wash with a CO2-independent media (Ref 18045088, ThermoFisher), cells were loaded with CCF2 (Ref K1032, ThermoFisher) for 1h30 at RT. Cells were then washed with CO2- independent media and the BlaM reaction was developed overnight at RT. Finally, the infected cells were washed once in PBS, fixed with formalin (Ref HT501128, Sigma-Aldrich) and analyzed using flow cytometry using the NovoExpress software. For experiments with GFP-LC3B and GFP-LC3B G120A transduced cells, a compensation was required and varied depending on levels of transduction. Thus, the entry levels were monitored using the value of cleaved-CCF2 rStaining using the following formula: rStain = (MFI CCF2-cleaved in infected cells - MFI CCF2-cleaved in uninfected cells) / (rSD MFI CCF2-cleaved in uninfected cells).

### Live-cell imaging

One day prior imaging, edited cells were seeded in an Ibidi 35mm high plate (Ref 81156, Clinisciences) and grew overnight at 37°C, 5% CO2. Then, the cells were pre-treated during 90 mintes with an anti-bleaching media (DMEM without phenol red, Ref 31053023, Gibco) containing 1% Penicillin-Streptomycin, 10% FBS and 1% Prolong Live Antifade Reagent (Ref P36975, ThermoFisher). Sir-Actin staining was performed during 15 minutes at 0.5µM and removed before infection. Cells were infected with viral particles for 30 minutes at 16°C for synchronization, then put 10 minutes in the slide chamber at 37°C to allow the following proper imaging. Movies were acquired using an inversed confocal microscope Nikon coupled with Spinning Disk Dragonfly Andor at an 40x objective. 488nm, 561nm and 637nm channels were acquired (GFP-LC3B, GagmCherry, Sir-Actin respectively) every 10s for 30 minutes by focusing on the top of the cells.

### Quantification of the distance between LC3B and the plasma membrane

Individual cells, selected using the Sir-Actin staining, were cropped using the Fiji software. Each cell was imported in the IMARIS software to quantify the distance between GFP-LC3B puncta and plasma membrane in 3 dimensions. Briefly, we identified the plasma membrane (Surface module) using the SirActin channel and applied a manual threshold with background subtraction and obtained a surface highlighting the cell contouring. Then, using the “Spots” module, we detected the GFP-LC3B puncta and we exported the measures “Shortest distance to Surface”, which indicate the distance of each GFP-LC3B puncta from the plasma membrane.

### Quantification of the GFP-LC3B puncta toward the viruses at the cell surface

The upper stacks (1.5µm of depth) were studied by generating a z-projection (based on maximum intensity). We chose blindly, using the GagmCherry and SirActin stainings, the incoming GagmCherry puncta at the plasma membrane that, lately, entered into the target cell. Tracking was performed manually by delimiting a zone of 1.5µm diameter around the viral particle, independently of its size, at each timeframe. Then, we counted the number of GFP-LC3B puncta in the delimited zone using the “Analyze particles” module after having applied a manual threshold determined for each cell and maintained throughout the timeframes.

### Statistical Analysis

Statistical analysis were performed using Graph Pad Prism software version 5. Differences were considered significant at *\*p<0.05, **p<0.01, ***p<0.001*. Specific statistical treatments are described in the figure legends.

## Acknowledgments

We thank Dr. Philippe Benaroch (Institut Curie INSERM U932, Paris) for the Env-mutated pNL4.3.Gag-mCherry (R5 tropic). We acknowledge the imaging facility MRI, member of the national infrastructure “France-BioImaging infrastructure” supported by the French National Research Agency (ANR-10-INBS-04, «Investments for the future»)”. This work was supported by the “ANRS- Maladies Infectieuses Emergentes », by SIDACTION, by the Centre National de la Recherche Scientifique (CNRS) and by the University of Montpellier.

## Funding

ANRS – Maladies Infectieuses Emergentes - grant number 244097 and 176196 (LE).

Sidaction – Grant 2022-1-FJC-13342 (BP).

Sidaction – Grant 2021-1-AEQ-12958 (RG).

ANR22 - CE17-0054-02 (MF).

Centre National de la Recherche Scientifique ; Université de Montpellier (LE, NC and RG).

