## Supplemental figures for "LC3B conjugation machinery promotes autophagy-independent HIV-1 entry in CD4+ T lymphocytes"

### Slide 1
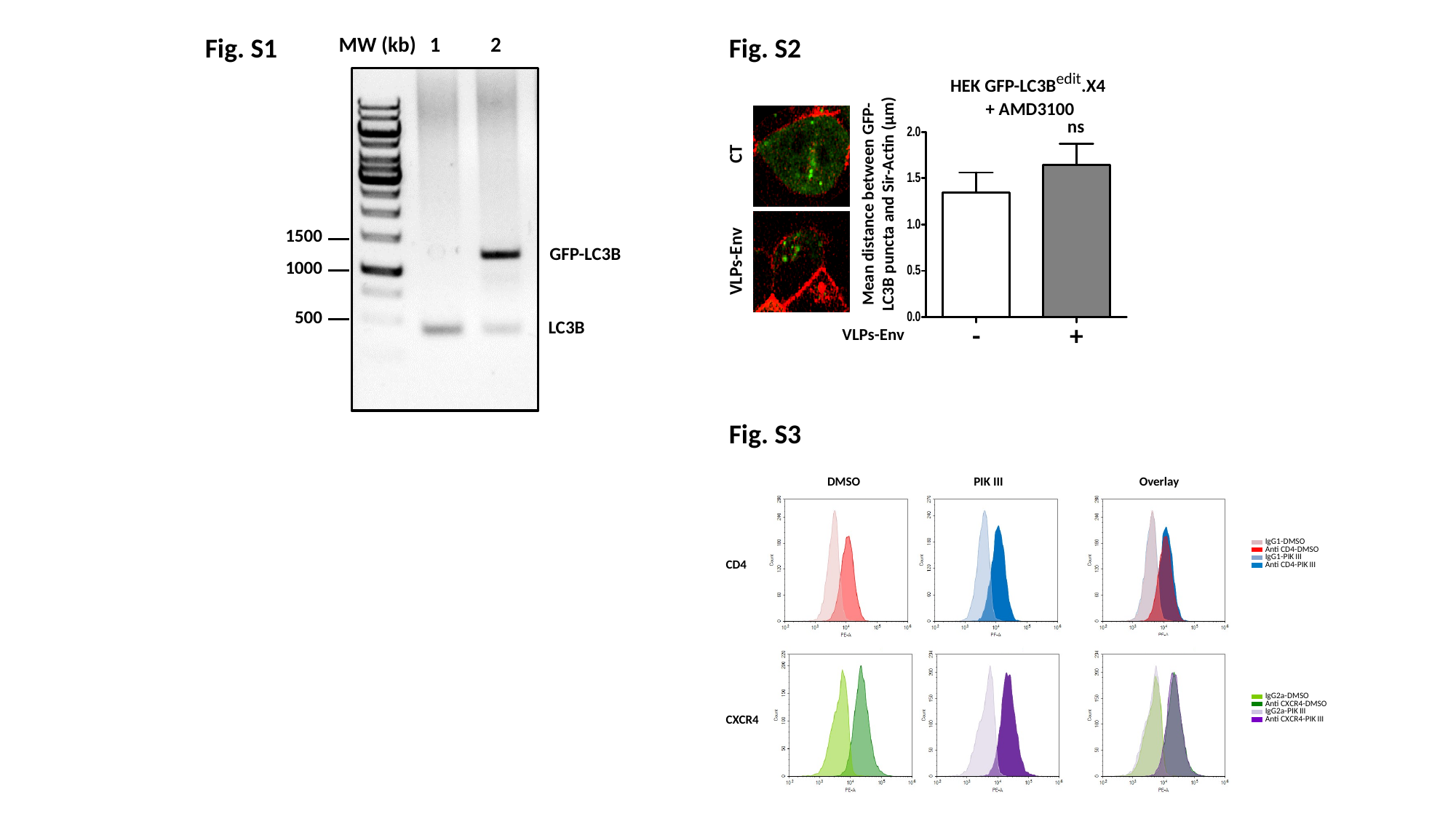

Fig. S1
MW (kb)
1
2
1500
1000
500
GFP-LC3B
LC3B
Fig. S2
HEK GFP-LC3B^edit^.X4
+ AMD3100
ns
CT
VLPs-Env
Mean distance between GFP-LC3B puncta and Sir-Actin (µm)
-
+
VLPs-Env
Fig. S3
